# SARS CoV-2 variant B.1.617.1 is highly pathogenic in hamsters than B.1 variant

**DOI:** 10.1101/2021.05.05.442760

**Authors:** Pragya D. Yadav, Sreelekshmy Mohandas, Anita M Shete, Dimpal A Nyayanit, Nivedita Gupta, Deepak Y. Patil, Gajanan N. Sapkal, Varsha Potdar, Manoj Kadam, Abhimanyu Kumar, Sanjay Kumar, Deepak Suryavanshi, Chandrashekhar S. Mote, Priya Abraham, Samiran Panda, Balram Bhargava

## Abstract

**Background:** The recent emergence of new SARS-CoV-2 lineage B.1.617 in India has been associated with a surge in the number of daily infections. This variant has combination of specific mutations L452R, E484Q and P681R reported to possibly enhance the transmissibility with likelihood of escaping the immunity. We investigated the viral load and pathogenic potential of B.1.617.1 in Syrian golden hamsters.

**Methods:** Two groups of Syrian golden hamsters (9 each) were inoculated intranasally with SARS CoV-2 isolates, B.1 (D614G) and B.1.617.1 respectively. The animals were monitored daily for the clinical signs and body weight. The necropsy of three hamsters each was performed on 3, 5- and 7-days post-infection (DPI). Throat swab (TS), nasal wash (NW) and organ samples (lungs, nasal turbinate, trachea) were collected and screened using SARS-CoV-2 specific Real-time RT-PCR.

**Results:** The hamsters infected with B.1.617.1 demonstrated increased body weight loss compared to B.1 variant. The highest viral load was observed in nasal turbinate and lung specimens of animals infected with B.1.167.1 on 3 DPI. Neutralizing antibody (NAb) and IgG response in hamsters of both the groups were observed from 5 and 7 DPI respectively. However, higher neutralizing antibody titers were observed against B.1.167.1. Gross pathology showed pronounced lung lesions and hemorrhage with B.1.671 compared to B.1.

**Conclusions:** B.1617.1 and B.1 variant varied greatly in their infectiousness, pathogenesis in hamster model. This study demonstrates higher pathogenicity in hamsters evident with reduced body weight, higher viral load in lungs and pronounced lung lesions as compared to B.1 variant.

**Summary:** B.1.617.1 is the new SARS-CoV-2 lineage that emerged in India. Maximal body weight loss and higher viral load in hamsters infected with B.1.617.1. It caused pronounced lung lesions in hamsters compared to B.1 variant which demonstrates the pathogenic potential of B.1.617.1.

## INTRODUCTION

Since, the first report of severe acute respiratory syndrome coronavirus-2 (SARS CoV-2) in Wuhan, China in 2019, the virus has constantly evolved leading to the emergence of new variants [1, 2]. The first variant of SARS CoV-2, D614G (B.1 lineage) became dominant and is prevalent worldwide since March 2020. Later mutations at different amino acid positions were reported with regional dominance [3] and were designated as variants under investigation (VUI). The variant of concerns (VOC) has also been identified in many countries with increased transmissibility, pathogenicity and phenomenon of immune escape.

Currently, multiple emerging variants have been circulating worldwide, of which, VOC B.1.17 (United Kingdom), Brazil P1, P2 and B.1.351 (South Africa) have seriously affected many countries and posed major public health challenges (PANGO lineages 2021). The emerging VOC of SARS CoV-2 variants B.1.1.7, B.1.351 and B.1.617 have been also detected and reported from India [4□6]. The second wave of the SARS CoV-2 has hit India hard and affected the population in large numbers. The number of new cases reported in India is increasing and averaging ∼ 0.4 million cases a day as of 30^th^ April 2021[7]. The state of Maharashtra shares 21% of the active cases (0.67 million) from among the reported cases of 3.17 million from the country [8]. The majority of these cases in Maharashtra have reported to harbor the SARS-CoV-2 B.1.617 lineage and its sub-lineages PangoLIN lineage [9].

B.1.617 variants have been associated with a surge of cases spreading rapidly not only in Maharashtra but have been identified in other states of India and abroad. B.1.617 variant has eight amino acid changes in the spike region. However, the evolution of multiple sub-lineages within B.1.617 lineage is a major concern. Sub-lineages B.1.617.1, B.1.617.2 and B.1.617.3 have a different set of unique substitutions and deletions [6, 9]. The origin of this variant is still unknown and as mentioned above, the presence of this variant has been identified not only in India but from 21 different countries [10]. Our recent study has shown that Covaxin (a whole virion inactivated vaccine) works against B.1.617 variant [6]. However, answering the question of this variant is the main cause of the surge of the second wave in India remains a challenge.

In an earlier study, we investigated the pathogenicity of the SARS CoV-2 B.1 and B.1.1.7 lineage in the Syrian hamster model [11]. It was observed that B.1 lineage of SARS CoV-2 variant produced interstitial pneumonia with marked alveolar damage and type-II pneumocyte hyperplasia in hamsters [11, 12]. In the present study, we investigated the viral load and pathogenicity of the B.1.617.1 variant in the Syrian hamster model and compared it with that of B.1 lineage.

## METHODS

### Ethics statement

The animal experiments were performed with the approval of the Institutional Animal Ethics committee and all the experiments were performed as per the guidelines of CPSCEA (NIV/IAEC/2021/MCL/01).

### Experimental design

Syrian golden hamsters of age 6-8 weeks were used. Nine hamsters each were inoculated intranasally with 0.1 ml of 104.5/ml TCID50 dose of SARS CoV-2 isolates i.e., B.1 (D614G) (GISAID number: EPI_ISL_420545) and B.1.617.1 (GISAID number: EPI_ISL_1669767) lineages propagated at ICMR-NIV, Pune as described earlier under isoflurane anesthesia [9,13]. Clinical signs and body weight of animals were observed daily and three hamsters each were euthanized on day 3, 5-and 7-days post-infection (DPI) to perform a necropsy. Throat swab (TS), nasal wash (NW) and organ samples (lungs, nasal turbinate, trachea) were collected following the necropsy.

### Real-time RT-PCR

Nasal wash and TS collected in 1ml viral transport medium and weighed organ samples (lungs, nasal turbinate and trachea) triturated in 1 ml media were used for RNA extraction using MagMAX™ Viral/Pathogen Nucleic Acid Isolation Kit as per the manufacturer’s instructions. Real-time RT-PCR was performed for the E gene for SARS-CoV-2 as well as for detection of sgRNA of the E gene using published primers [14, 15].

### Histopathology

Lungs samples collected during necropsy were fixed in 10% neutral buffered formalin. The tissues were processed by routine histopathological techniques for hematoxylin and eosin staining. The lung lesions were graded from 0 to +4 as nil, minimal, mild, moderate and severe based on the vascular changes, inflammatory changes and bronchial/ alveolar damage.

### Plaque reduction neutralization test

The assay was performed against both the isolates as described earlier [16]. Four-fold serial dilution of hamster serum samples mixed with an equal amount of virus suspension and incubated at 37°C for 1 hour. Further 0.1 ml of the mixture was inoculated in a 24-well tissue culture plate containing a confluent monolayer of Vero CCL-81 cells and was incubated at 37°C for 60 min and overlay medium was added to the cell monolayer, which was further incubated at 37°C in 5% CO2 incubator for 4-5 days and PRNT50 titres were calculated.

### Virus titration

The lungs and nasal turbinate samples were used for virus titration in Vero CCL-81 cells. A hundred microliter of the sample was added onto a 24**-**well plate with Vero CCL-81 monolayers and incubated for one hour at 37°C. The media was removed and the cell monolayer was washed with PBS and was incubated with maintenance media with 2% FBS in a CO2 incubator. The plate was examined daily for any cytopathic effects (CPE) and the culture supernatant from the wells showing CPE was confirmed by real**-**time RT**-**PCR.

### Statistical analysis

Graphpad Prism version 8.4.3 software was used to assess the statistical significance. A non-parametric two-tailed Mann–Whitney test was used to compare between two groups and p-values of < 0.05 were considered to be statistically significant.

## RESULTS

### B.1.617.1 infection lead to higher weight loss in hamsters compared to B.1

Weight loss was observed in a higher percentage of hamsters infected with B.1.617.1 but it was not significant in comparison with the B.1 infected hamsters (Figure 1). It was observed that both the B.1 and B.1.617.1 infected hamster groups demonstrated a decline in the body weight starting from 3^rd^ day post-infection (DPI). The mean value for the percent change in bodyweight of B.1 infected hamsters was -0.63± 3 (95% CI: -2.96 to 1.70; N=9) at 3 DPI. The mean body weight decreased further by 2.9 (N=6) and 0.9 (N=3) fold on the 5th and 7th DPI in comparison to the 3rd DPI. However, a maximum change in body weight was observed for B.1.617.1 infected hamsters (mean ± SD: -2.66± 3.2, N=9) with 95% CI:-5.14 to -0.18 on 3 DPI. The body weight was further decreased 3.5 fold at the 5 DPI and 6.4 fold at 7 DPI in comparison to the 3 DPI. This indicates that B.1.617.1 affected the hamster’s health more in comparison to B.1 variant.

**Figure 1:**
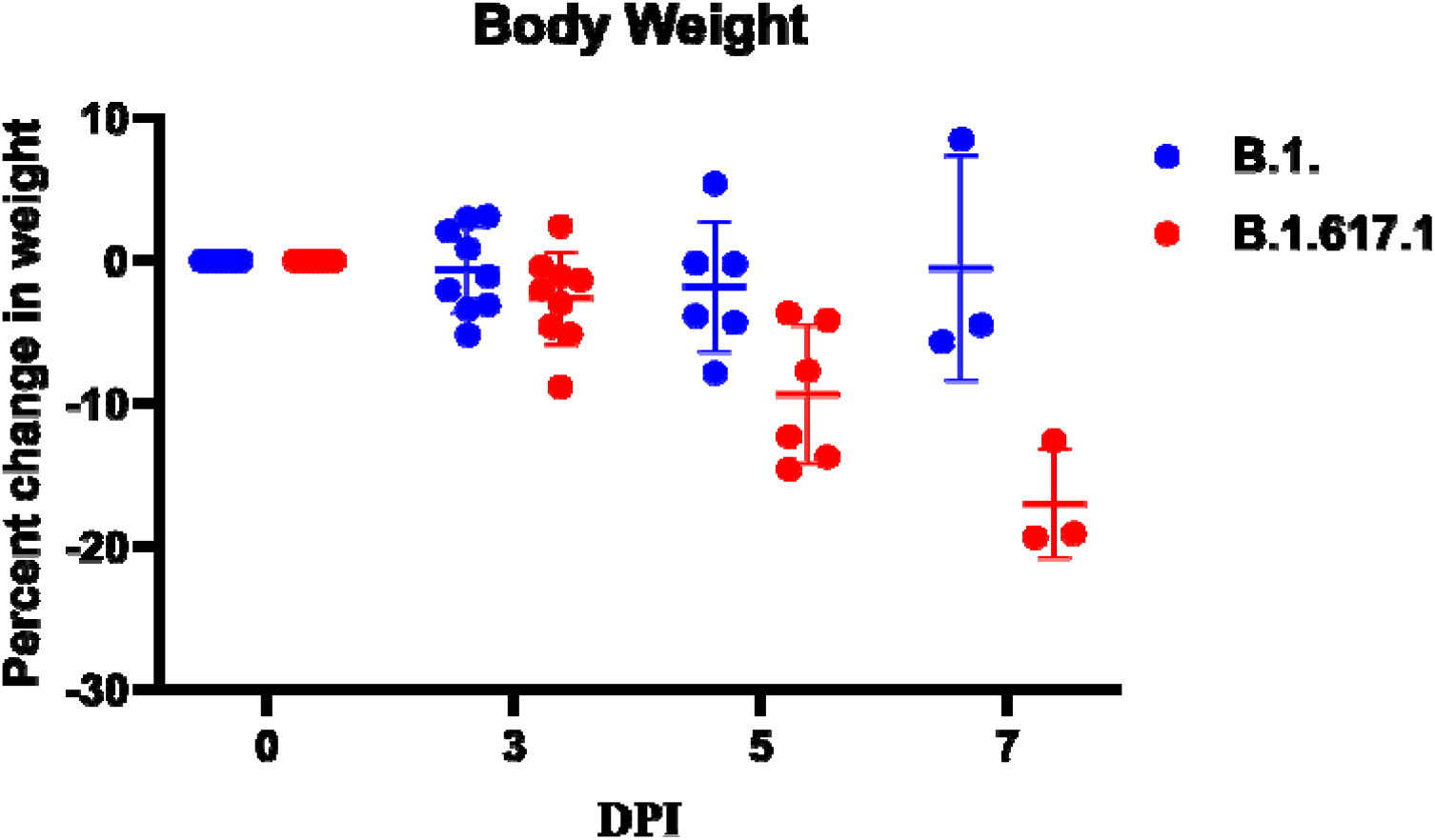
Percent body weight change in the hamsters infected with SARS-CoV-2: The body weight change in hamsters following exposure among hamsters to SARS CoV-2 variants B.1 and B.1.617.1. The statistical significance of difference in weight loss between the SARS CoV-2 variants B.1 and B.1.617.1 was assessed using the two-tailed Mann–Whitney test between two groups; P values of < 0.05 were considered to be statistically significant. Mean and standard deviation is depicted for animals in each group.

### Genomic and sub-genomic SARS-CoV-2 viral loads in hamster infected with B.1 and B.1.617

The viral loads in the throat swab (TS), nasal wash (NW) and organ samples were found comparable and did not show any statistical significance between the two variants. Sub-genomic RNA (sgRNA) could be detected in the samples up to 5^th^ DPI. The highest viral load of genomic RNA (gRNA) was observed in the nasal turbinate’s for B.1 and lungs for B.1.617.1 on 3^rd^ DPI, although the difference was not statistically significant with the B.1 variant. The viral load showed a decreasing trend on 7^th^ DPI in all the samples of hamsters infected with both the variants (Figure 2 A-J).

**Figure 2:**
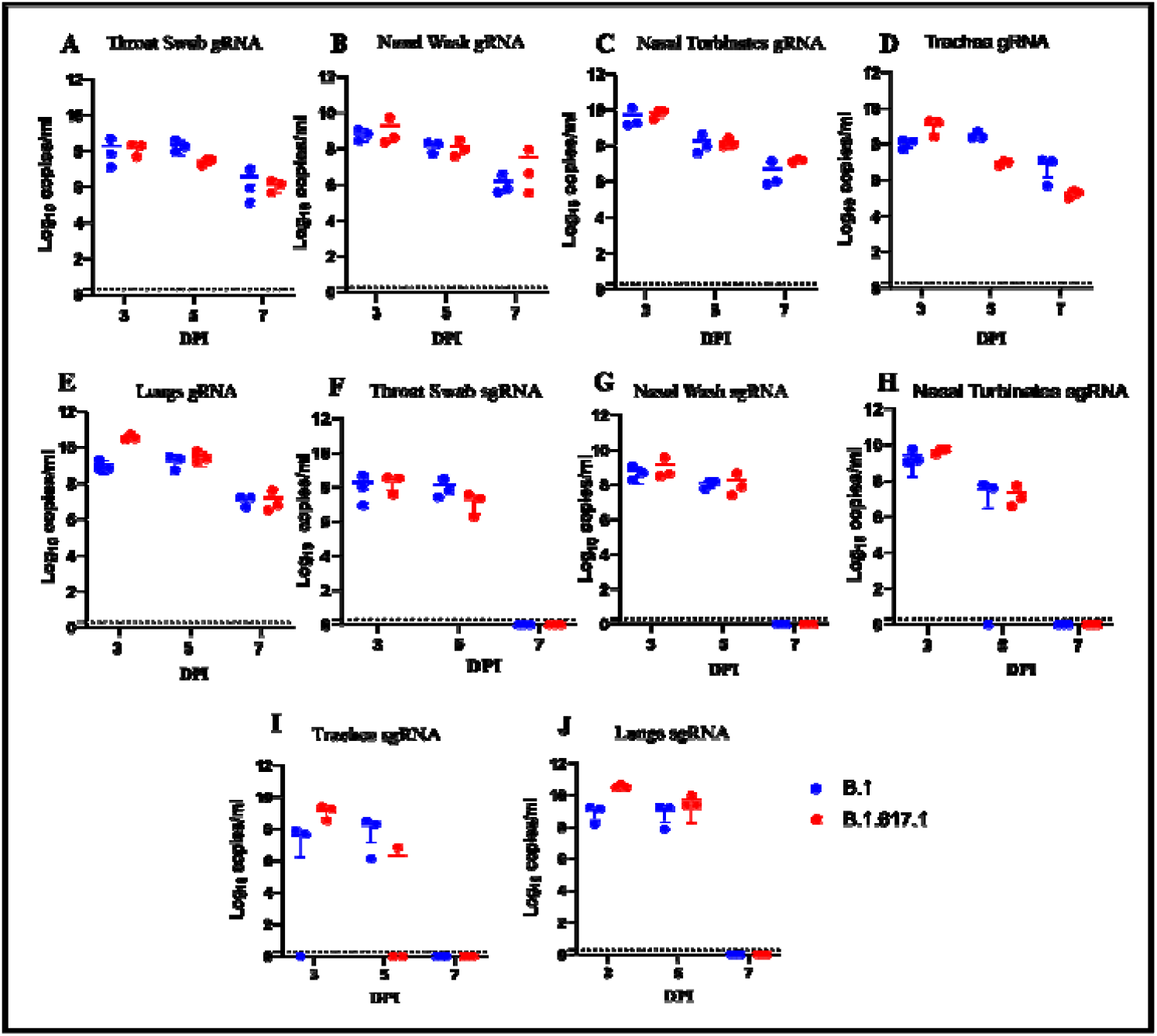
Scatter plot for the SARS-CoV-2 genomic and sub genomic viral loads in hamsters using real time RT-PCR: Genomic viral RNA titers in the swab and organ samples of hamsters infected with SARS CoV-2 variants B.1 and B.1.617.1. **A)** Throat swab **B)** Nasal wash **C)** Nasal turbinates **D)** Trachea **E)** Lungs. Sub-Genomic viral RNA load in **F)** Throat swab **G)** Nasal wash **H)** Nasal turbinates **I)** Trachea **J)** Lungs. The statistical significance between the SARS CoV-2 variants B.1 and B.1.617.1 was assessed using the two-tailed Mann–Whitney test between two groups; P values of < 0.05 were considered to be statistically significant. The dotted line on the figures indicates the limit of detection of the assay. Mean and standard deviation is depicted for three animals in each group.

### Antibody response in hamsters post infection

Neutralizing antibody (NAb) response in hamsters was observed from 5^th^ DPI against B.1 and B.617.1 variants in infected hamsters whereas the anti-SARS CoV-2 IgG antibodies could be detected by ELISA at 7^th^ DPI except in one of the hamsters infected with B.1 (Figure 3A). The variation of antibody response was found to be not significant between the groups both by ELISA and neutralization. The neutralizing antibody geometric mean titer (GMT) for the B.1.617.1 infected hamsters against B.1 was 260(100.7-672.7) while against B.1.617.1 GMT was 424 (110.4-1632) on 5 DPI. However, on 7^th^ DPI, there was a 0.77-fold reduction against B.1 while a two-fold increase against B.1.617.1. B.1 infected hamsters have reduced NAb titer against the B.1.617 in comparison to B.1 starting from the 5^th^ DPI (Figure 3B). A two-fold increase in the NAb titer was observed against the B.1 on 7 DPI. However, on 7^th^ DPI, there was a six-fold increase against B.1.617.1. However, higher NAb titers for the B.1.617.1 infected variant were observed than for the B.1 infected variant on 7 DPI.

**Figure 3:**
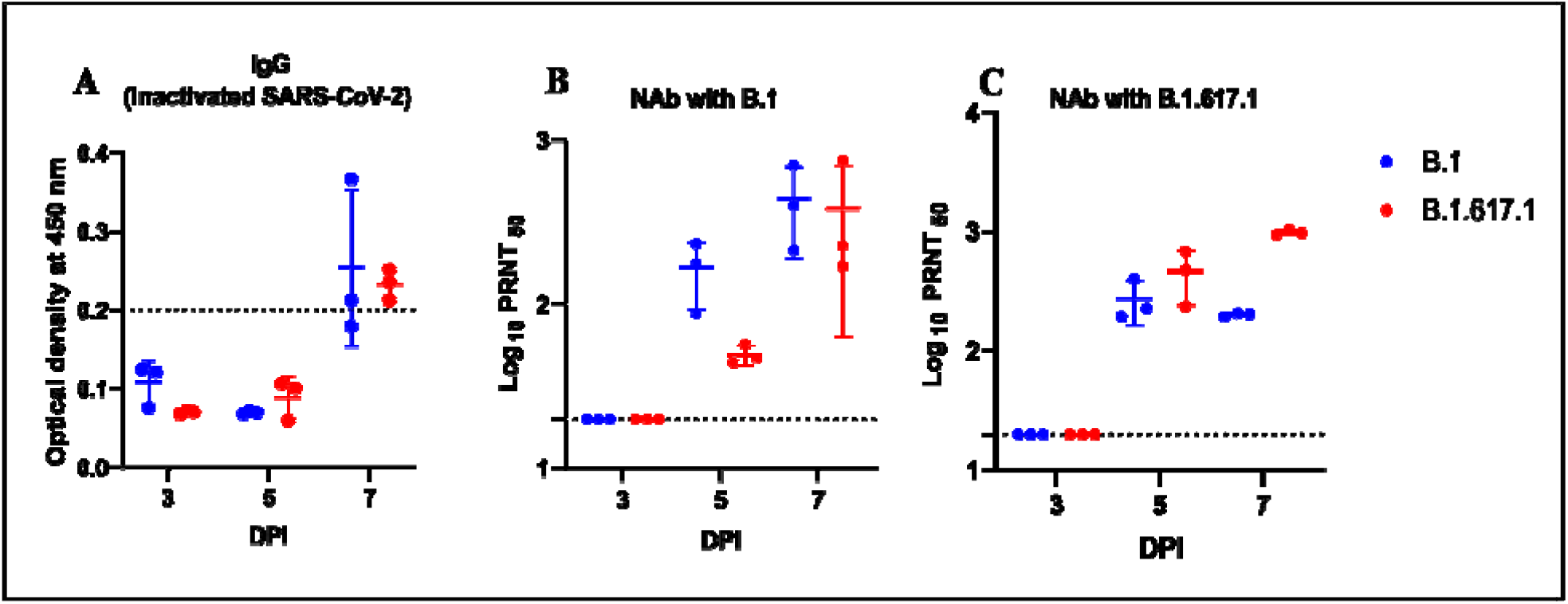
Anti-SARS-CoV-2 antibody response: **A)** IgG response using anti-SARS CoV-2 IgG ELISA in animals infected with B.1 and B.1.617 variants on 3, 5 and 7 DPI. **B)** The neutralizing antibody response of the hamster infected with B.1 and B.1.617 variants against B.1. **C)** The neutralizing antibody response of the hamster infected with B.1 and B.1.617 variants against B.1.617.1. The statistical significance between the SARS CoV-2 variants B.1 and B.1.617.1 was assessed using the two-tailed Mann-Whitney test between two groups; P values of < 0.05 were considered to be statistically significant. The dotted line on the figures indicates the limit of detection of the assay. Mean and standard deviation is depicted for three animals in each group.

### B.1.617.1 induced more severe lung lesions in hamsters

Gross examination of lung specimens showed pronounced congestion and hemorrhages on 5^th^ and 7 DPI in the case of the B.1.617.1 as compared with the B.1. The lung lesions in hamsters infected with the B.1 variant ranged from minimal to mild whereas the B.1.617.1 variant developed moderate lesions (Table 1). In the case of the B.1 variant, the pneumonic changes observed were minimal to mild which included inflammatory cell infiltration, focal consolidation and mild congestion (Fig 4A-4C). The pronounced changes (moderate to severe) with mononuclear infiltration in the alveolar interstitial space, interstitial septal thickening, consolidation and pneumocyte hyperplasia were observed with B.1.617.1 variant consistently till 7DPI (Fig 4D-4F).

**Table 1.** Final grade of involvement of hamsters lungs infected with B.1.617.1 and B.1 (D614G) variant

| **Days post**<br>**infection** | **B.1 (D614G)** |  |  | **B.1.617.1** |  |  |
| --- | --- | --- | --- | --- | --- | --- |
|  | **Animal-1** | **Animal-2** | **Animal-3** | **Animal-1** | **Animal-2** | **Animal-3** |
| **3** | +1 | +2 | +2 | +3 | +2 | +2 |
| **5** | +2 | +2 | +2 | +3 | +3 | +3 |
| **7** | +2 | +2 | +2 | +3 | +3 | +3 |

**Figure 4:**
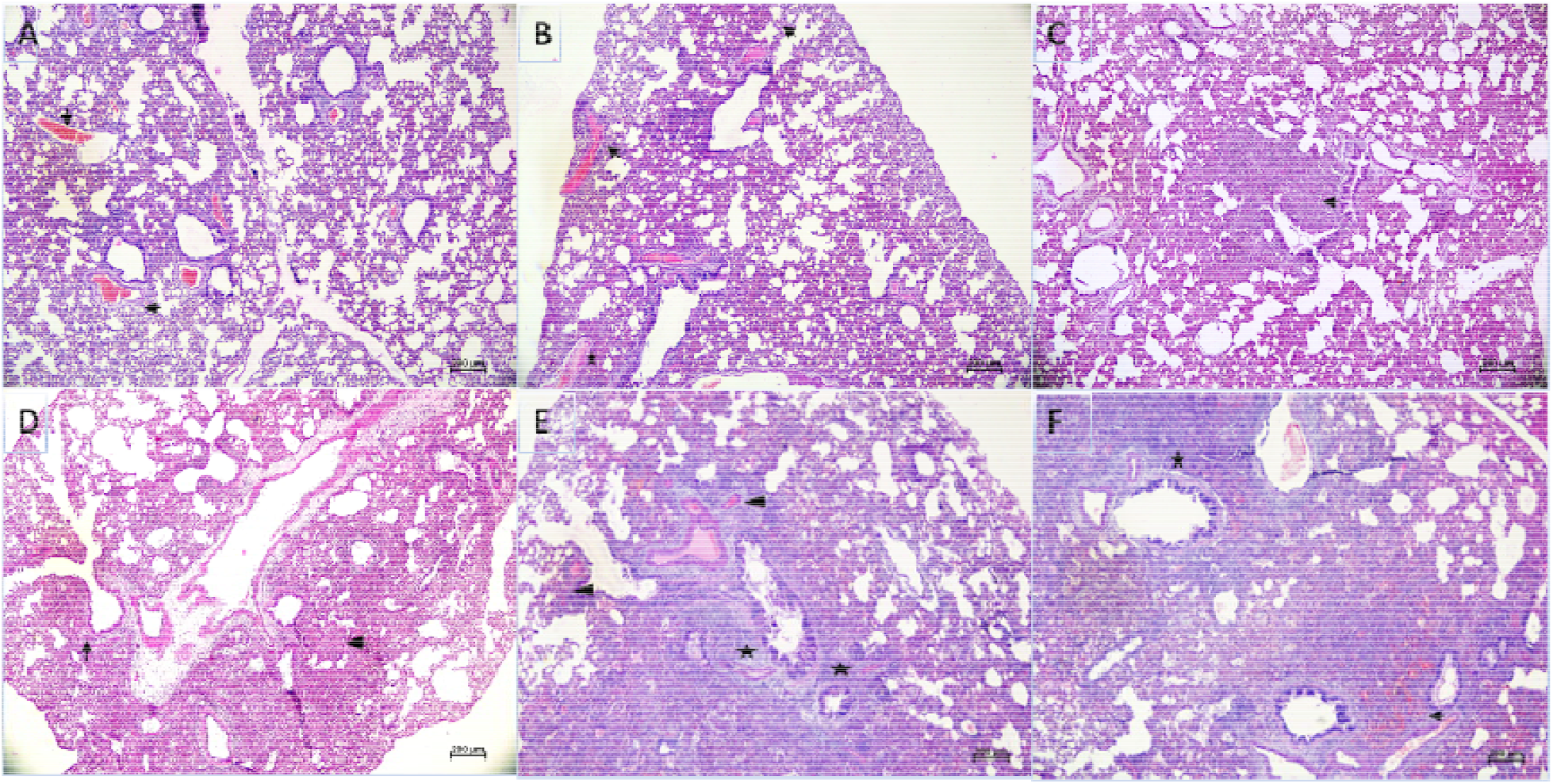
Histopathological changes in hamster’s lungs infected with SARS-CoV-2 variants: Lung section of hamsters infected with B.1 showing **A)** congested vessels (arrow) on 3DPI **B)** marked congestion (arrow), hemorrhages (star) and focal areas of inflammatory cell infiltration on 5 DPI **C)** peribronchial mononuclear infiltration (arrow) and areas of septal thickening on 7DPI **D)** Lungs section of hamsters infected with the B.1.617.1 showing mutifocal areas of inflammatory cell infiltration (arrow) on 3 DPI **E)** congestion (arrow),peribronchial infiltration (star) and alveolar septal thickening on 5 DPI **F)** areas of hemorrhages (arrow), inflammatory cell infiltration (star) and consolidative changes on 7 DPI.

## DISCUSSION

The emergence of a new lineage of SARS-CoV-2 (B.1.617.1) has raised concerns among public health experts. A total of 791 SARS-CoV-2 sequences of B.1.617 and its sub-lineages were reported from India till 30 April 2021 and deposited in the Global Initiative on Sharing All Influenza Data (GISAID) database (https://www.gisaid.org/). The predominant lineage being the and B.1.617.3, and these are characterized by the presence of L452R, E484Q, and P681R amino acid changes, while the B.1.617.2 lacks the E484Q mutation. The presence of L452R, E484Q, and P681R is linked with increased hACE2 receptor binding affinity and possible NAb escape [15, 16].

The massive surge of cases in India necessitated the study of pathogenicity of the variant in the hamster model. Using B.1.617.1 virus isolate, we demonstrated the pathogenic potential of this variant in the hamster model in comparison to B.1. Syrian hamsters are a widely used animal model for SARS-CoV-2 and the clinical signs/ or endpoints of infection are weight loss and viral RNA/infectious virus titer in target organs [17]. The increased severity of B.1.617.1 infection in hamsters was demonstrated by the higher bodyweight reduction and lung viral load, and more severe lung histopathological changes as compared to that of B.1. Although the viral load and the bodyweight loss were not statistically different, the severity of the lung lesions produced by B.1.617.1 indicates increased pathogenicity of this isolate as compared to B.1. The increased receptor binding affinity attributed to the characteristic mutations of the B.1.617.1 isolate might have contributed to the increased disease severity. Although the B.1 variant is known to enhance the virus infectivity and transmissibility, it has been reported not to cause severe disease in humans [18, 19]. Hou et al. have also demonstrated that the B.1 does not enhance the pathogenicity in hamster models [20]. Besides this, Abdelnabi et al. reported a very efficient infection of B.1.1.7 and B.1.351 variants in the respiratory tract of hamsters [21].

The protective efficacy of the immunity generated from an initial SARS-CoV-2 infection against subsequent reinfection is puzzling. Studies in animal models have shown that prior infection results in protective immunity [22]. In the present study, we have evaluated the neutralization efficacy of the sera of hamsters infected with B.1.617 against B.1 variant and vice versa and found robust neutralizing immune response in both conditions. Re-infection studies need to be performed to understand the extent of the protection by these neutralizing antibodies in decreasing the viral load and severity of the disease. We recently reported that sera from BBV152 vaccine recipients and recovered cases of COVID-19 produced robust neutralizing antibodies against B.1.617 variant [6]. This means that the immunization program has great relevance against the different variants circuiting in India and other country and vaccines will have the potential to protect the recipient against severe morbidity following infection.

Overall in this study, Syrian hamsters infected with B.1.617.1 SARS-CoV-2 variants showed higher viral load and greater loss of body weight but it was not significant in comparison with that caused by the variant of B.1 linage. However, the increased pathogenicity of the B.1.617.1 isolate was evident by the severity of the lung histopathological changes in hamsters. BBV152 vaccine recipients can neutralize the new variant, further bolstering the need for rapid vaccine roll-out.

## Funding

Financial support was provided by the Indian Council of Medical Research (ICMR), New Delhi at ICMR-National Institute of Virology, Pune under intramural funding of ‘COVID-19’.

### Competing interests

No conflict of interest exists.

## Acknowledgments

Authors gratefully acknowledge the team members of Maximum Containment Facility, ICMR-NIV, Pune including Mr. Prasad Sarkale, Mr. Shreekant Bardkar, Ms. Pranita Gawande, Mrs. Ashwini Waghmare, Ms. Kaumudi Kalele, Ms. Jyoti Yemul and Ms. Manisha Dhudhmal for providing excellent technical support.

## Notes

### Competing Interest Statement

The authors have declared no competing interest.

